# Combinatorial analysis of gene regulatory network reveals the causal genetic basis of breast cancer and gene-specific personalized drug treatments

**DOI:** 10.1101/2020.04.10.035998

**Authors:** Mohammad Kawsar Sharif Siam, Mohammad Umer Sharif Shohan, Easin Uddin Syed

**Author notes:** Corresponding Author: Mohammad Kawsar Sharif Siam, Computational Molecular Biology and Bioinformatics Lab, Department of Pharmacy, BRAC University, 41 Pacific Tower, Mohakhali, Dhaka 1212, Bangladesh.

## Abstract

Cancer is the major burden of diseases around the world. The incidence and mortality rate of cancers is mounting up with the passage of days. Breast cancer is the most demoralizing cause of death, where both diseases interlocked with each other due to some genetic, biological and behavioral motives. The molecular mechanism of breast cancer through which they crop up and manifest together remains questionable. The genetic basis of protein-protein interactions and gene networks has elucidated a group of gene regulatory systems in Breast cancer. Thus, the extraction of all genomic and proteomic data has enabled unprecedented views of gene-protein co-expression, co-regulation, and interactions in the biological system. This study explored the biological system to develop a gene-disease interaction model by implementing the extracted genomic and proteomic data of Breast cancer. The disease-specific and correlated genes were pulled out and their cabling studied by PPI, disease pathway and drug-disease interaction data to articulate their role in disease development. By analyzing mined genes that are related to breast cancer, a network model is also proposed. Exploration of all the correlated genes, Hub and common genes have given some promising pieces of evidence surrounding the genetic networking models. The result of this prospective study disclosed breast cancer mediated crosslinking or possible metastatic relation on a genetic basis. Moreover, other diseases like prostate, colorectal and ovarian cancers are at the same risk and might count into consideration. The finding provides a narrative broad approach for understanding the genetic basis of these fatal diseases by the pathway analysis with gene regulatory network evaluation.

## Introduction

As the world is experiencing growth in the number and lifespan, the burden of cancer will inevitably increase.^1^ World Health Organization (WHO) identified cancer as one of the four leading threats to human health and development. It is estimated there will be 22.2 million new cases of cancers diagnosed annually worldwide by 2030.^2^ It has been predicted that in 2018 there will be 18.1 million new cancer incidence and 9.6 million deaths from cancer worldwide. The prevalence of breast cancer (11.6% of all cases) is the second leading cause of cancer death (6.6% of all cancer deaths).^3^

Breast cancer (BC) is the most common type of non-skin cancer in women and the second most common cause of cancer death.^3^ BC (predominantly affecting females) refers to a malignant tumor that has developed from breast cells (usually in the epithelial lining of the ducts or lobules). The genetic basis behind breast cancer involves the generation of altered cells, owing to a mix of deleterious single nucleotide polymorphisms (SNPs) and non-inherited somatic changes (mutations, deletions, translocation, etc.). These cells give rise to the characteristics of cancer such as unrestricted proliferation, and metastasis.^4^

Although several genes have linked to the tumorigenesis of BC, the molecular mechanism of BC tumorigenesis is still unclear. Therefore, it is essential to identify additional relevant genes that can be used as candidates. These genes can then be used as a biomarker for early diagnosis and clinical research. It is laborious and expensive to determine these genes by wet-lab experiments alone. A computational approach to identify has been successful in recent years due to its ability to debug the complex biological network.^5^ Finding candidate genes will help us understand the genetic factors of breast cancer in women, develop new treatments, and better personalization of treatment.^6^

Various integrative approaches have been developed to create a identify candidate genes for complex diseases such as BC. In general, there are two ways candidate genes can be identified. The “Candidate gene” approach identifies a correlation between genetic variants and the disease. It also thought to be more economical. The “genome-wide scanning” involves scanning the whole genome, mostly using human linkage data. However, this approach is more expensive than the previous approach. Both approaches may result in producing hundreds of candidate genes. Too many candidate genes lead to confusion amongst researchers trying to validate them in the wet lab. Thankfully, bioinformatic approaches can be adopted to analyze the candidate genes using data from protein-protein interactions, gene regulatory networks, SNP, and expression data.^5^

In this study, an approach was taken to identify common BC related genes through comprehensive text mining. Along with data exploration, genes associated with BC were explored. All the disease-related data were further analyzed by using various types of bioinformatics tools and resources to identify key candidate hub genes correlated with one another in BC. This could help oncologists in understanding the gene not only individually but as a network in cancer, which can be exploited for better treatment.

## Methods

This present work has developed gene interaction network models comprising of high throughput genomic and proteomics data for BC. The approach is illustrated in (Figure 1). This study aimed to identify the role of BC genes and their regulation.

**Figure 1:**
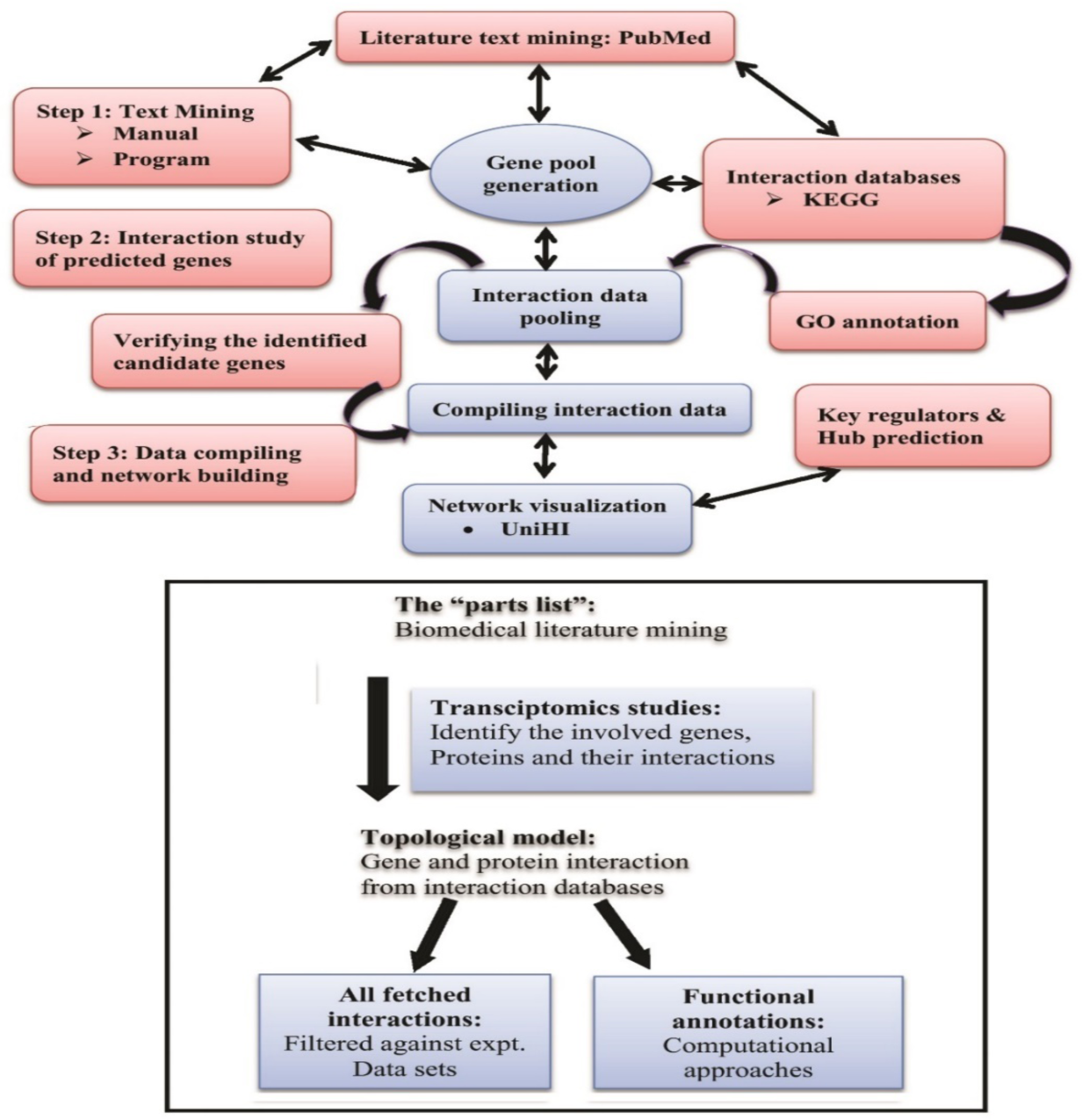
Collated strategy and concept for unified network development of breast cancer in three successive stages using biochemical literature mining, topological model generation and fetched interaction data along with fine computational approaches.

### Development of the network model

The network model has been developed in three stages. The first stage consisted of the “parts list” generation (Figure 1) using biochemical literature mining. The second stage was the topological model generation by combining all the pooled genes and their interactions from interactive databases. In the third and final stage, the fetched interaction data were filtered against experimentally validated interaction (e.g. yeast two-hybrid, chip-chip interactions, immune co-precipitation, and array co-expression) datasets using computational approaches. By identifying the genes, the pathways they are involved in and the interactions they are part of, using a systems approach; the integrated network of BC has been generated considering all experimentally verified PPI data of the regulated genes. Finally, from data, “Hub genes” and from literature reviewing, “common genes” were identified.

#### Stage 1: Primary disease-oriented gene pool generation through extensive data mining

PubMed (http://www.ncbi.nlm.nih.gov.pubmed) is one of the centers of scientific publications collection, which contains biomedical articles from Medline and life sciences journals. Citations include links to full-text articles from PubMed Central (PMC). PMC is the U.S. National Institute of Health (NIH) free digital archive of biomedical and life sciences journal literature serving the critical role of providing access to publish literature toward the first step in synthesis and translation of genomic research easily and comprehensively. PubMed supports keyword-based searching and information retrieval (IR). To increase the accuracy and efficiency for discovering the relationships between important biological entities, the present study combined manual (literature search) and automated (computational) tools to identify the genes and gene products by computationally capturing the related knowledge embedded in textual data.

#### Stage 2: Construction of protein-protein interaction (PPI)

PPI maps have a considerable impact on the discovery and synthesis of molecular networks. Thus, generating human protein interaction maps has become an important tool in biomedical research for the elucidation of molecular mechanisms and the identification of new modulators of disease processes. There have been several web-based tools available for biologists. The Unified Human Interactome Databases (UniHI, http://www.unihi.org) provides a comprehensive platform to query and access human protein-protein interaction (PPI) data.^7^ UniHI has integrated PPI resources to provide a comprehensive platform for querying the human protein interaction. The idea was not to replace a single database but to design convenient portal access to human protein interaction data for the biomedical research community.^7^ Additionally, it facilitates the identification of network topologies, which would not be detectable if PPI resources examined separately. Pathway information can provide useful clues about the function and the dynamic of the interactions. Especially for the elucidation of network structure. UniHI provides the possibility to examine the intersection of canonical pathways from the Kyoto Encyclopedia of Genes and Genomes (KEGG) with the extracted networks.^8^ UniHI scanner does not only show the proteins included in the pathway but also the KEGG annotations of the interaction between nodes. In this way, it enables researchers to detect possible modifiers of pathways as well as proteins involved in the cross-interactions between pathways.

#### Stage 3: Filtration and validation of PPI data

Advances in recent genome-wide interaction projects have generated a wealth of PPI data. To understand the complexity, it is essential to extract meaningful information in the context of the biological system. This necessitates not only identification of the functions of individual proteins, but also the validation of the physical interactions and biological processes in which they participate. The emergence of large-scale protein-protein interaction maps has opened up new possibilities for systematically surveying and studying the underlying biological system which recently integrates protein interaction data with pathway data from the human gene expression atlas.^9^ For normalization of intersections, however, only the common protein interactions were used. In UniHI, maps are subsequently clustered based on the interaction overlap. For all comparisons, it is notably larger than zero, which is the expected value for comparison of random maps. Using UniHI, users can filter interacting proteins by using a minimum expression threshold, and the PPI network retrieved from UniHI can be simplified to include either highly or poorly expressed proteins.^7^ In this study interaction data was filtered from already available experimentally validated datasets.

## Results and Discussion

### Comprehensive data mining and generation of gene pooling

Candidate genes responsible for BC were pooled through text mining (Table 1). Candidate genes were selected based on several criteria’s given below:

**Table 1:**
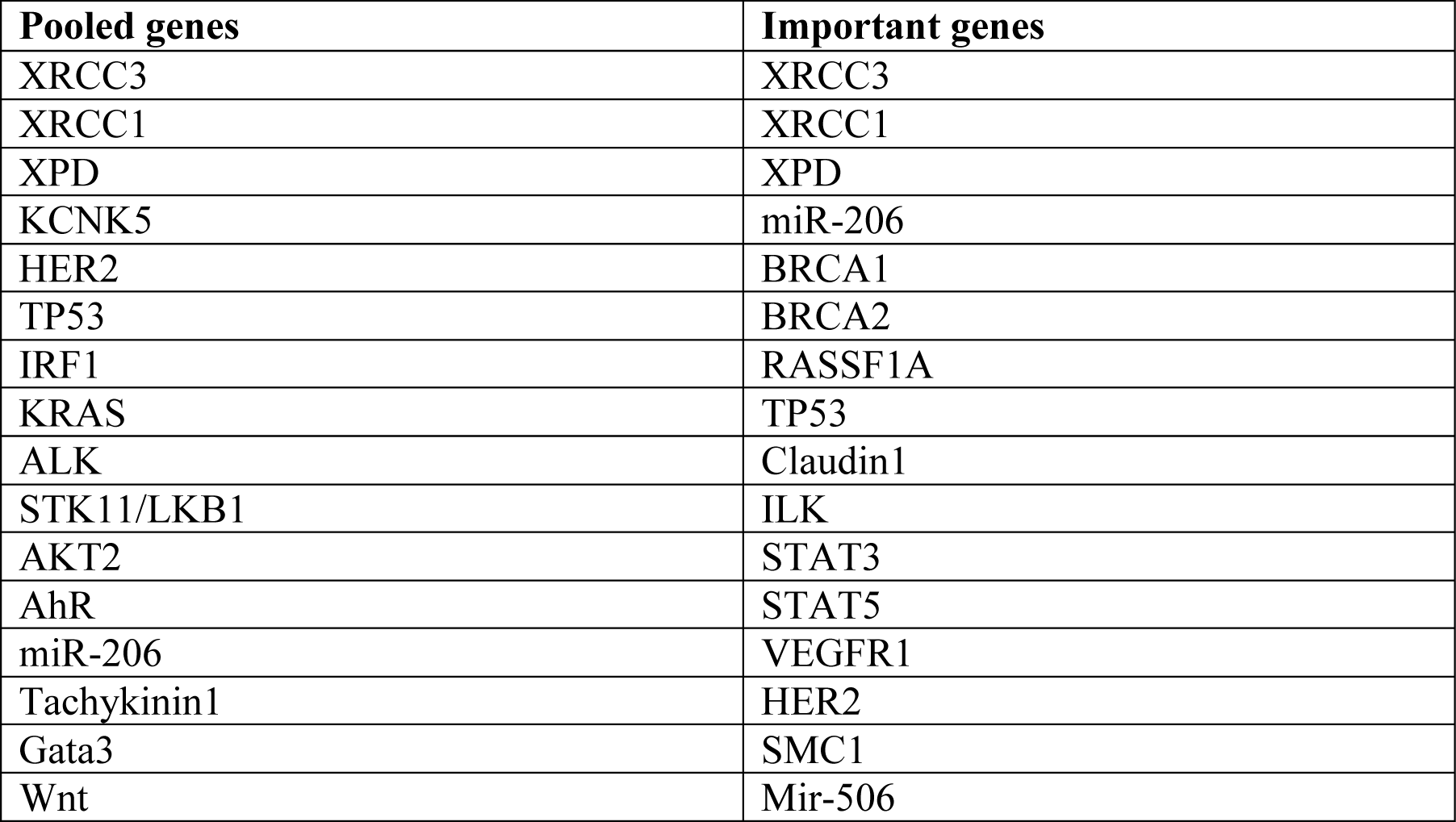
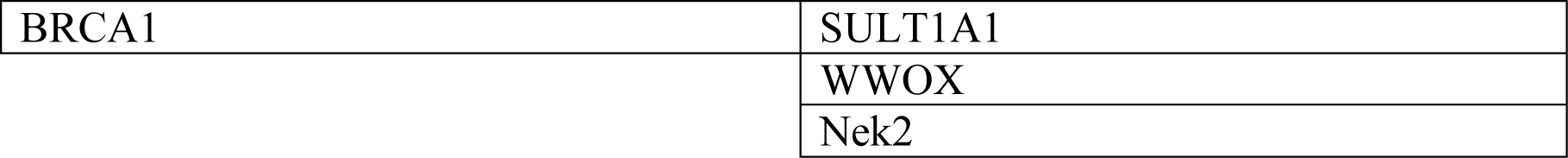
36 Pooled genes and important genes for breast cancer.

- Genes experimentally were proven to be associated with any of the above-mentioned diseases,
- Genes experimentally were proven to be associated with two or more concerning diseases,
- Genes predicted computationally to be associated with an individual disease,
- Genes predicted computationally to be associated with two or more concerning diseases and
- Genes are found in the KEGG pathways database to be associated with the individual disease.

The next step of gene extraction was the generation of important genes. KEGG pathways and literature citations were used to identify the role of genes in BC. These were labeled as important genes in this study. 36 BC related genes were labeled important (Table 1).

### Finding Protein-Protein Interaction networks (PPIs)

Genes usually do not act as individual units; they collaborate in overlapping pathways. This co-regulation is important in the pathogenesis of several diseases. A simple enrichment analysis has been applied to characterize cancer on the network level including pathways and protein-protein interactions. The computational approach used in this study was based on functional links derived from co-expression and co-regulation profiles.

The gene interaction data used to build the network were based on direct physical interactions that are either experimentally derived or computationally predicted. To integrate pathway information and to derive cellular network information of the selected genes, functional annotations from pathway databases such as KEGG, and protein-protein interaction data was added.^10,11^

To facilitated the findings, UniHI (Unified Human Interactome)^7^ and Pathway commons^11^ (Figure 2) which are web-based PPI tools, were used. The present work attempted to find significantly interconnected sub-networks.

**Figure 2:**
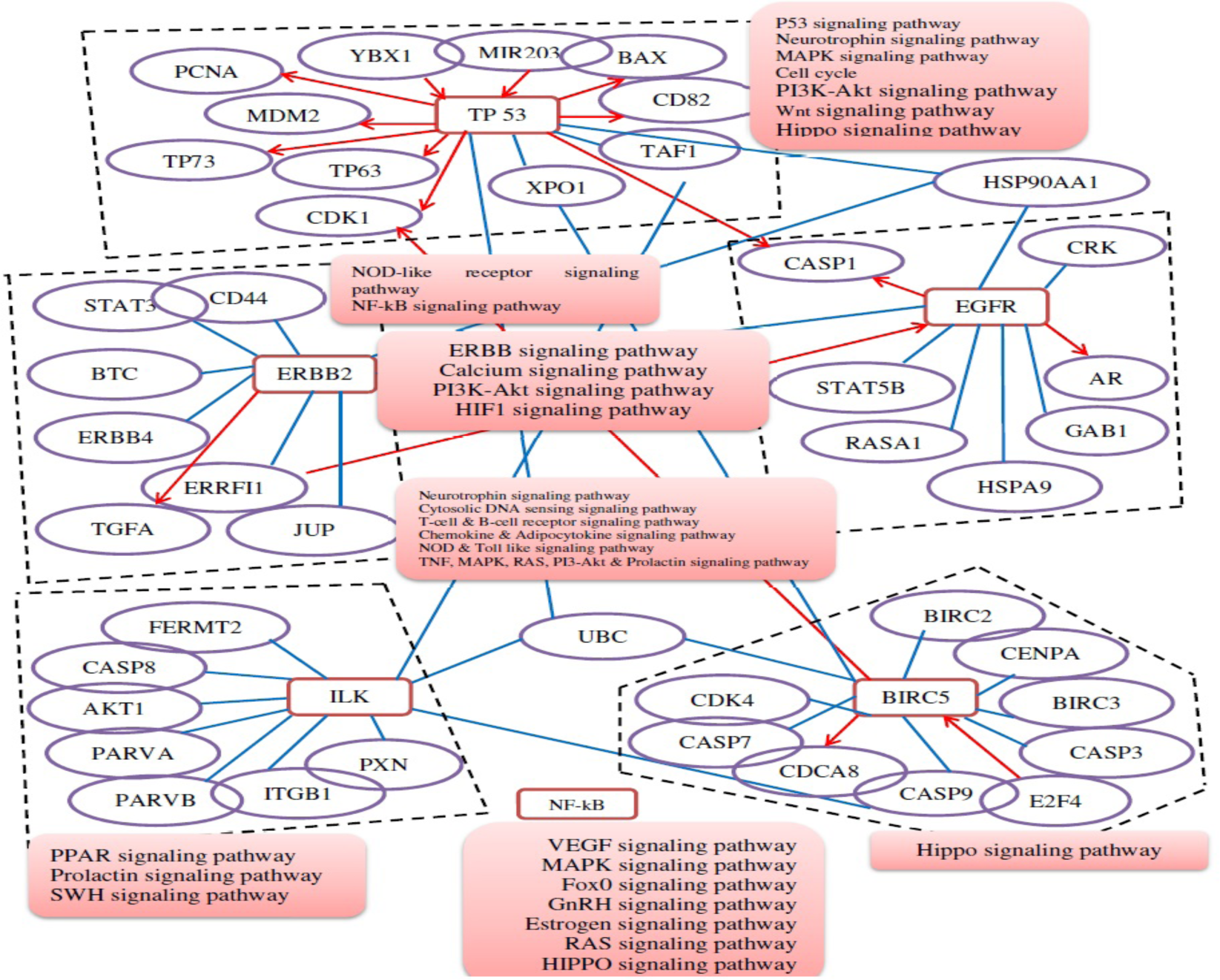
Protein-protein network revealed common genes and their direct or indirect actions in breast cancer with the association of some concomitant signaling pathways. Most of genes had many similar signaling pathways to exert their genetic characteristics.

### Data compilation and network visualization

Network subgraphs can be network modules, motif clusters or other network neighborhoods. A network module can be defined as a subgraph consisting of highly interconnected nodes that may fulfill a particular biological function.^12-14^ At this stage of network building, the pooled important genes (Table 1) for various diseases were integrated into network models and visualized using UniHI or Pathway commons or Cytoscape.

The models have been generated through several stages of screening in subsequent steps. The pooled data have been filtered, verified for authentication and most importantly validated with experimentally derived data to generate meaningful and interpretable output graphs for different diseases. Therefore, a BC network was established, containing a set of proteins that are associated with the disease. The hub nodes form the backbone of a network are considered an important measurement for the similarity between protein interaction networks.

To propose a hypothetical model of BC development and to develop credible treatments, genes were considered based on their regulation, association and regulatory pathways. The information was gathered from the extensive literature review and the small broken pieces of puzzles were stitched by bioinformatics tools to develop a hypothetical model for the diseases. The network was then further analyzed to find the Achilles heel for both diseases together which can be targeted to mitigate both diseases in one shot. The complete regulatory pathways are yet to be confirmed and it is still unclear how a risk factor influences the regulatory network of the healthy cell to be converted into a cancerous cell. The generated regulatory networks associated with the extracted genes via pathways like HER2 or ErbB2 Signaling Pathway, NF-kappa B Signaling Pathway, HIF-1 Signaling Pathway, PI3K-Akt Signaling Pathway, MAPK Signaling Pathway, NGF Signaling Pathway, and P53 Signaling Pathways has been implicated in various types of cancer.^15-23^

The Growth of a cell depends on the transduced signal by a signaling molecule that is sent to the neighboring cells. When the signal from the extracellular signaling molecule reaches the nucleus of a cell, cell proliferation mechanisms start, and all the machinery required for cell division are transcribed and translated by the cell itself. A wrong or defective signaling molecule may trigger an abnormal signaling pathway and ultimately leads to cancer. ^24^

In the detailed hypothetical model of BC development (Figure 3, it was found that cancer prerequisite phenomena to cancer such as DNA damage, abnormal DNA synthesis abnormal protein synthesis are triggered by the activation of the common pathways involving HER2/ErbB2, EGFR, TP53, NF-kB, surviving or BIRC5, ILK and VEGF genes (Figure 3). These genes were also found to be over-expressed in the cancer cells, which might be another reason to produce an excess amount of oncoprotein leading the cell into carcinogenesis.^25^

**Figure 3:**
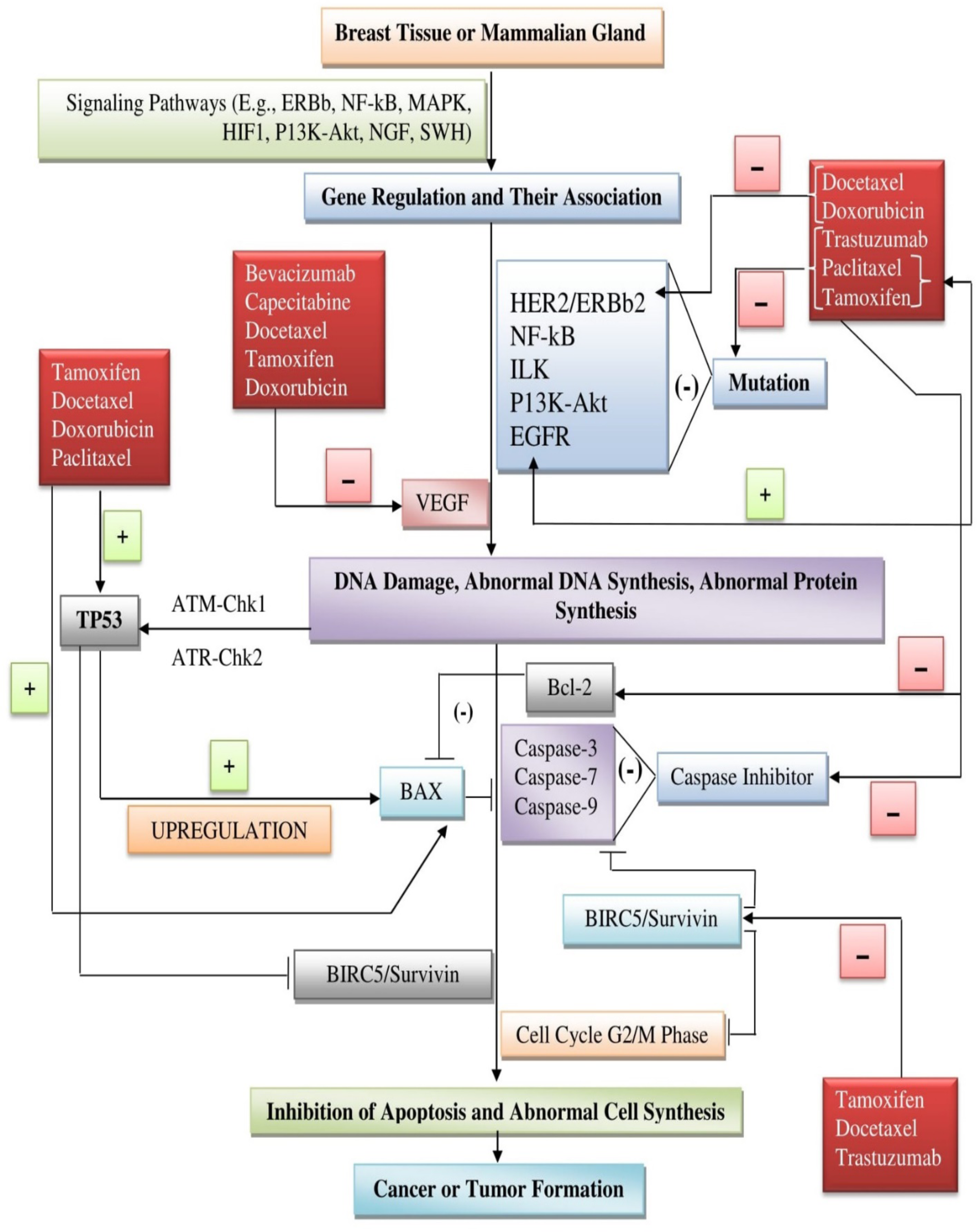
Proposed hypothetical model of developing breast and lung cancer & credible drug treatment, according to the validated and scrutinized gene networking and interconnecting data.

DNA damage can be sensed by the ATM and ATR kinases and in response to the damaged DNA, they activate Tumor Protein p53 (TP53) via ATM-Chk1 and ATR-Chk2 signaling pathways which further repairs damaged DNA, arrests cell cycle, programmed cell death and several other downstream processes (Figure 3).^26,27^ TP53 initiates apoptosis by activating Bax gene which in turn forms Bax-Bax homodimer and initiates apoptosis. This process can be disrupted in cancer cells by Bcl-2 protein which forms Bax-Bcl2 heterodimer and gives cell a survival signal.^28^ Apoptosis is also important in destroying cells which has been metastasized. Normally, metastatic cancer cells undergo genetic changes that allow them to avoid apoptosis and travel to another place. To take control over the cell the apoptotic process recruits and activates Procaspase enzymes which further activates executioner apoptosis factors called caspases like Caspase-3, Caspase-7, Caspase-9 resulting in the death of the metastatic cancer cell (Figure 3). Though the role of executioner caspases is questionable for their non-apoptotic roles in cancer cells, they are found to be not mutated in the cancer tissues in human and trigger proliferation and damage repair.^29^

To counterbalance the fatal effect of caspases and to maintain the cellular homeostasis, cell deploys an apoptosis regulatory network by activating Bcl-2 gene. Bcl-2 acts by cleaving the caspases and prevent apoptosis in healthy cells.^30^ The level of expression of these proteins in the cell are strictly regulated by the cell. However, the altered level of Bcl-2 and caspases are observed in various breast cancer cell lines.^31,32^ For surviving in the apoptotic situation, cancer cells not only depend on the Bcl-2 gene but also triggers the synthesis of ‘Survivin’, an inhibitor of apoptosis (IAP) protein encoded by the BIRC5 gene. Survivin can inhibit apoptosis by both caspase-dependent and independent pathways.^33^ Apart from its function of avoiding apoptosis, it also helps in spindle apparatus assembly in the G2/M phase of the cell cycle.^34^ TP53 can inhibit survivin synthesis in the cell transcriptionally by down-regulating survivin mRNA synthesis.^35^ Even though these predictions need further experimental validations, the hypothetical model proposed in this study could be a starting point to understand and visualize the gene expression network in BC development. Numerous drugs are now available for the treatment of breast cancer. But all types of drugs are not directed towards all types of breast cancer. A drug, which is productive to an individual, maybe a contraindication, less effective or ineffective for others. From that point of understanding, it can notably be said that after understanding the genotype of the breast cancer treatment pattern should be fixed and that could be the standard treatment.

Breast cancer associated with the mutation of genes can be treated using multiple drugs. Tamoxifen and Doxorubicin are used for treating breast cancers associated with the mutation of protein encoded gene HER2 or ErbB2. Along with these drugs Docetaxel, Paclitaxel and Trastuzumab are used for correcting their expression. Breast cancer-associated with EGFR mutation treated with drugs like paclitaxel and tamoxifen. VEGF decreases susceptibility and downregulates expression in the case of breast cancer under treatment with Bevacizumab, Capecitabine, Docetaxel, Tamoxifen and Doxorubicin drugs.^36^

Oncogenes those are liable for any type of DNA damage or abnormal synthesis of DNA and protein upregulates tumor suppressor protein TP53. Drugs like dictate, paclitaxel and doxorubicin play sound role to increase the expression of TP53, apoptosis regulator Bax for the inhibition of the carcinogenic process by triggering the apoptosis in cancerous cells.^36^ On the other hand, Tamoxifen acts by downregulating the expression of the Bcl-2 level in the cell and it does not alter the TP53 and Bax expression level.^37^

Apoptosis suppressing gene Bcl-2 can be treated with Docetaxel, Paclitaxel, Tamoxifen, and Doxorubicin (Figure XX). These drugs inhibit the reaction of Bcl-2 and decrease their expression in the cell which clarifies increase expression of protein family-like Bax promoting the apoptosis process. These classes of drugs also play a positive role additionally during preventing the inhibition of apoptotic factors like Caspases. As such Tamoxifen, Docetaxel and Trastuzumab inhibit BIRC5 expression and promote the expression of apoptosis factors of caspases and normalize the cell cycle process and promote regular cell division and repairing of DNA (Figure 3). Tamoxifen decreases BIRC5 mRNA expression.^36^

## Conclusion

It can be hypothesized from the generated model that BC may develop from the interactions of highly interconnected gene network and the credible drug treatment can be given at the molecular level, after evaluating the genotype of the BC. From this study of BC, some novel findings provided a useful approach for understanding the genetic basis, networking, co-regulation and pathways of the diseases. The integrated network model provides useful functional linkages not only among the groups of genes but also diseases. It is anticipated that derived models would be of great benefit to a wide research audience, including those who are involved in the identification of disease biomarkers and drug development.

## Acknowledgment

This research received no external funding.

## Declaration Of Conflicting Interests

The author(s) declared no potential conflicts of interest with respect to the research, authorship, and/or publication of this article.

## Author Contributions

Mohammad Kawsar Sharif Siam conceived, designed and supervised the experiments, performed the experiments, analyzed the data, contributed materials/analysis tools, prepared figures and/or tables, authored or reviewed drafts of the paper, approved the final draft.

Mohammad Umer Sharif Shohan conceived and designed the experiments, performed the experiments, analyzed the data, authored or reviewed drafts of the paper, approved the final draft.

Easin Uddin Syed analyzed the data, authored or reviewed drafts of the paper, approved the final draft.

## Notes

### Competing Interest Statement

The authors have declared no competing interest.

